# Excitation energy transfer between higher excited states of photosynthetic pigments: 1. Carotenoids facilitate B → Q band conversion in chlorophylls

**DOI:** 10.1101/2023.01.26.525634

**Authors:** Jan P. Götze, Heiko Lokstein

## Abstract

Chlorophylls (Chls) are known for fast, sub-picosecond internal conversion (IC) from ultraviolet/blue absorbing (“B” or “Soret” states) to the energetically lower, red light-absorbing Q states. Consequently, excitation energy transfer (EET) in photosynthetic pigment-protein complexes involving the B states has so far not been considered. We present, for the first time, a theoretical framework for the existence of B-B EET in tightly coupled Chl aggregates, such as photosynthetic pigment-protein complexes. We show that according to a simple Förster resonance energy transport (FRET) scheme, unmodulated B-B EET likely poses an existential threat, in particular the photochemical reaction centers (RCs). This insight leads to so-far undescribed roles for carotenoids (Crts, this article) and Chl *b* (next article in this series) of possibly primary importance.

Here we show that B → Q IC is assisted by the symmetry-allowed Crt state (S_2_) by using the plant antenna complex CP29 as a model: The sequence is B → S_2_ (Crt, unrelaxed) →S_2_ (Crt, relaxed) → Q. This sequence has the advantage of preventing ~ 39% of Chl-Chl B-B EET, since the Crt S_2_ state is a highly efficient FRET acceptor. The likelihood of CP29 to forward potentially harmful B excitations towards the photosynthetic reaction center (RC) is thus reduced. In contrast to the B band of Chls, most Crt energy donation is energetically located near the Q band, which allows for 74/80% backdonation (from lutein/violaxanthin) to Chls. Neoxanthin, on the other hand, likely donates in the B band region of Chl *b*, with 76% efficiency. The latter is discussed in more detail in the next article in this series. Crts thus do not only act in their currently proposed photoprotective roles, but also as a crucial building block for any system that could otherwise deliver harmful “blue” excitations to the RCs.

## Introduction

Carotenoids (Crts) are pigments found in many organisms, with various functions.^1^ In photosynthesis, Crts act as (i) structure stabilizing components and accessory light-harvesting pigments, (ii) chlorophyll (Chl) triplet excited state quenchers, and (iii) possibly, enabling photoprotection via non-photochemical quenching (NPQ). NPQ is also assumed to originate from other mechanisms.^2–6^ An example for a typical spatial arrangement of Chls and Crts in light-harvesting pigment-protein complexes (LHCs) is given in Figure 1A.

**Figure 1:**
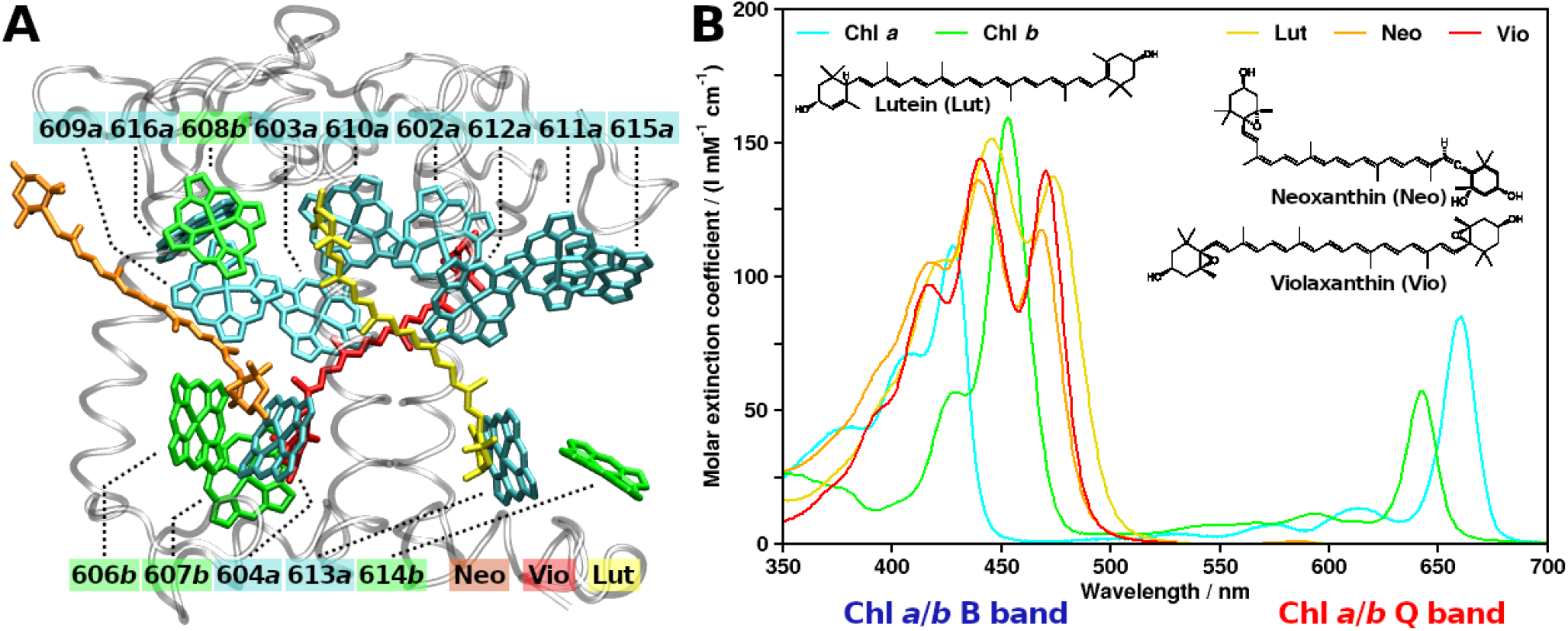
Structural arrangement and spectroscopy of typical plant photosynthetic pigments. A: CP29 antenna complex from Pisum sativum, PDB structure 5XNL by Su et al. (2017). Viewpoint along Thylakoid membrane plane, looking towards the center of the photosystem II supercomplex. Stromal side at the top, lumenal side at the bottom. Protein grey and transparent, Chls a shown as cyan, Chls b as green (Chl numbering as given in earlier CP29 structures). Crts also shown, lutein in yellow, neoxanthin in orange and violaxanthin in red. B: Ultraviolet/visible region absorption spectra of CP29 chromophores (see text and Table 1 for sources).

Despite the importance of Crts, a crucial aspect of Crt photophysics has, so far, not been highlighted: The spectral region in which Crts absorb correlates with the Chls B band absorption (Figure 1B). Chls and Crts exhibit strong absorption in the blue-green spectral region (300 nm and longer wavelengths). Especially for Chl *a* and *b*, in plant systems, the overlap between the spectra is high. For an accessory, light harvesting pigment, one would rather expect a shifted absorption to cover more of the Chls’ “green gap” (cf., *e.g.*, the phycobiliproteins). Instead, Crts in most organisms containing Chl *a/b* exhibit strong absorption and significant overlap with the higher energy Chl bands. Notably, the Chl B band consists of multiple states^7^. For the sake of simplicity, they are treated as a single entity in the following.

In this article, the “competition” between Chls and Crts is explored on the basis of incoherent excitation energy transfer (EET). To keep matters simple, only Förster resonance energy transfer (FRET) theory will be employed.^8,9^ In the context of Crts, FRET has the explicit downside that it relies on a point dipole approximation (PDA). For Chls, PDA holds much better than for Crts, since Chls are comparatively compact rings, while Crt chromophores are extended chains.^10^ The pigments’ “centers” are needed to assign a distance between the transition dipole moments (TDM) for FRET rate calculations (see below). This is thus much less defined for Crts than for Chls. Center of mass (COM) distances might in both cases be valid at large separations, but at short distances, PDA based on the COM coordinates becomes less valid as well. Still, these shortcomings can be managed through exploring the distance limit, and future studies may include more advanced coupling schemes, covering the full Coulomb coupling, such as TrESP^11^ or transition density cube (TDC).^10,12^ In fact, such a study is currently being conducted in our lab. For the proof of concept, FRET suffices, since predicted rates by multipole methods (*e.g*., TrESP or TDA) should be generally on the same order of magnitude as the rates predicted by FRET.^10^ FRET further has the distinct advantage of being easily comparable with experimental data. Coherent transport for strongly coupled chromophore pairs is also possible but will not be explored here; its occurrence would however only strengthen our arguments.

The aforementioned spectral competition between Chls and Crts implies the existence of potentially harmful, Chl photophysics beyond IC, *i.e.*, B-B excitation energy transfer (EET). As such, the first task of this article is to show that such interactions exist. This is easily done by considering the results by Leupold *et al*.,^13^ who have experimentally shown the corresponding Chl *a/b* B band emission properties. Further, although unmentioned by the authors, Zheng *et al*. have shown in a theoretical study that B-B EET precedes any B → Q events in a Chl-Chl dimer model (B-B EET lifetime < 10 fs, as seen in Figure 3 of their publication).^14^ The results shown below in the present article paint a similar picture, for the first time providing a physical basis and a reason for Crts to prevent B-B EET. Note that the detrimental effects of blue irradiation to the photosynthetic reaction center (RC) are known, with multiple mechanistic proposals.^15–17^ It is however photophysically of no concern if the high-energy quanta reach the RC via direct irradiation or EET, the effect should be the same.

This article will further investigate the potential of Crt back-donation to Chls - purely on the basis of the “bright” (allowed) Crt state. It has been known for a long time that the vibrational relaxation (VR) of the strongly absorbing Crt state (S_2_) is very fast (sub-100 fs^18^) and the actual reorganization energy of this process is large for such small changes in molecular geometry.^7,19,20^ Despite detectable S_2_ emissions of certain Crts, EET processes in plants between Crts and Chls have, to our knowledge, not been attributed to FRET (although shown for bacteriochlorophylls almost 30 years ago^21^). It is well-established that most of the (weak) fluorescence of Crts arises from S_2_, not S_1_.^1^ Crts thus intrinsically violate Kasha’s rule^22,23^ and assuming that the S_2_ state is a FRET donor seems to reasonable.

Consequently, this article will explore the following: First, the Chl-Chl interactions will be modeled. We will explicitly include all Chl bands, showing that the long-standing neglect of B-B excitation EET may have resulted in a misconception of the Crt role. Second, we will investigate the effect of introducing Chl → Crt EET as a competing element to Chl-Chl interactions. Third, the inverse process (Crt → Chl EET) after Crt VR will also be analyzed.

## Methods

### FRET model

To obtain the FRET rates (in ps^-1^) between a donor D and an acceptor A^8,9,24^

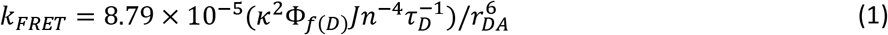

several excited state/pigment-specific and conformation-dependent parameters need to be known: *κ* or *κ*^2^ is the orientation factor, with *κ*^2^ ranging from 0 to 4, with the value 4 representing perfect alignment of the involved TDMs and chromophores. Φ_*f*(*D*)_ is the fluorescence yield of the donor state, *J* (in M^-1^ cm^-1^ nm^4^) is the spectral overlap, for which a variety of different definitions exist (see Eq. (2)).^24,25^ *n* is the refractive index of the medium (assuming a protein environment,^26^ a value of 1.4 is used here) and *τ_D_* the lifetime of the donor state (in ps^-1^). Finally, *r_DA_* is the distance (in Å) between the acceptor and donor compounds using the PDA. The Förster radius R0 is the distance *r_DA_* at which 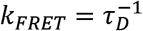. As already noted above, defining *r_DA_* between spatially extended compounds such as Crts is not straightforward, and the values computed here for very close pigment pairs must be used with caution.^10^ Comparing two compounds of similar character on the basis of FRET is justifiable, as the model can be built such that the actual *r_AD_* is not relevant (see below).*J* can be obtained^9,24^ from

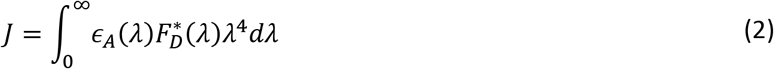

in units of M^-1^ cm^-1^ nm^4^, with *ϵ_A_*(*λ*) as the molar extinction spectrum of the acceptor (in M^-1^ cm^-1^), the wavelength *λ* (in nm) and the normalized emission spectrum of the donor 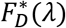(in nm^-1^). For the case presented here, the Chl overall absorption and Q band emission spectra by Li^27^ in diethylether are used for Chl *a* and *b;* the B band emission of these compounds is taken from Leupold *et al*.^13^ When restricting the Chl absorbance to either the Q or B band, the bands were separated at 470.5/495 nm for Chl *a*/*b*. For the spectra of Crts, the corresponding sources are listed in Table 1 contains three Crts present in the CP29 complex (Figure 1A; lutein (Lut, yellow), neoxanthin (Neo, orange) and violaxanthin (Vio, red)). Further, β-carotene (Bcr) is an ubiquitous Crt found in peripheral antennae as well as the RCs^35^ and zeaxanthin (Zea) is a replacement for Vio under light stress.^20,34^ Finally, peridinin (Per) is the primary light harvesting pigment in the peridinin-chlorophyll a-protein (PCP).^2–4,36,37^ PCP only contains Per and Chl *a*, with Chls *a* acting as the connection to the RCs. As such, Per should provide an improved donor capability to Chl *a*, allowing us to validate our model.^2,3,32,38,39^

**Table 1:**
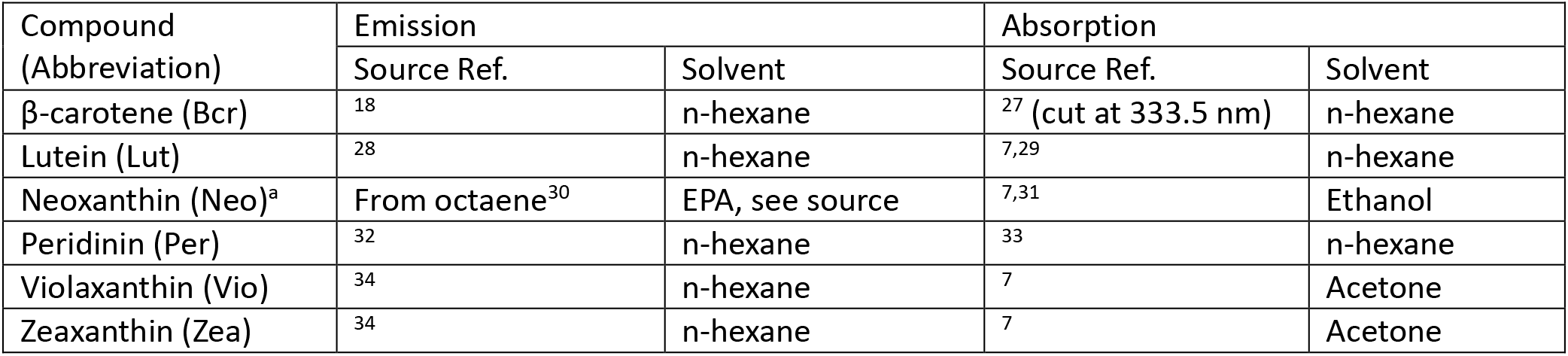
Literature sources of Crt spectra used in this article, with the corresponding solvent. ^a^: Neoxanthin emission unavailable, using octaene instead.

Finally, a FRET network will be set up, with acceptor Chls and Crts competing for excitation energy. For every donor, the total FRET efficiency is computed as

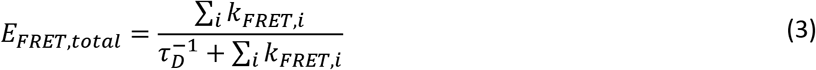

over all acceptors *i*. Note that *τ_D_* is considered invariant regardless of the presence of the acceptors.

### Spectral parameters

The main advantage of FRET is that it provides a direct connection between the experimentally measured spectral properties of the involved compounds and the EET rate. For the Chl Q band, the required values are well known, and we take a Q band yield of Chl *a/b* of 0.32/0.117 from our own work.REF??? The Q band lifetimes of 6.3/3.2 ns were taken from the literature.^40^ The most challenging parameters in Eq. (1) are those that relate to the spectral properties of the higher-energy B states (Chls) and S_2_ (Crts) as donors. For those, Φ*_f_* and *τ_D_* are known to be small and obtaining exact values requires careful measurements. In the case of Crts, corresponding values could only be found for Bcr in diethylether^18^ (1.50 10^-4^ for Φ_*f*(*Bcr,S*_2_)_ and 163 fs for *τ*_*Bcr,S*_2__); however the variance between Crts in the exact values is small.^1^ For the Chl Soret emission/internal conversion (IC), corresponding spectra and values are also only sparsely available.^13,14^ While the *τ_D_* values for Chl *a/b* could be found as 100/58 fs,^41^ the measurements by Leupold *et al*. only allow for an estimate of Φ_*f*(*Chl,B*)_ (“less than 10^-4^”).^13^ It is thus prudent to validate these values from other means: a theoretical approach.

It has been known for more than 50 years that the extinction spectrum can be reliably related to the rate of fluorescence via the corresponding Einstein coefficients, if the shape of the fluorescence spectrum is known.^42,43^ Then, the expression to obtain the rate of fluorescence, *i.e.*, the Einstein coefficient for spontaneous emission, is

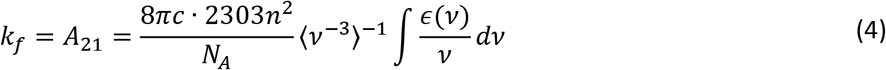

with *c* being the speed of light and *N_A_* as Avogadro s number. *v* is chosen here instead of *λ* for unit compatibility (wavenumbers). The resulting rate *k_f_* is in the time units chosen for *c*. Brackets indicate an intensity-weighted average of the fluorescence spectrum. For Eq. (4), the absorbance of the B band was set to end at *v_max_* 29700/25450 cm^-1^ for Chl *a/b;* a maximum value is not required for other equations, as those consider overlaps to emission. The fluorescence yield Φ*_f_* can then be obtained via its definition

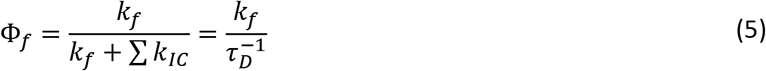

### Structural model

The CP29 complex (Figure 1A) will be used as an example, the corresponding site energies^44^ and distance matrices can be found in the supporting information (SI, Tables S1, S2 and S3). The site energies have been included as a shift of the Chl *a* or *b* spectra before entering in the calculation of *J* (Eq. (2)), while the distances are used in Eq. (1) directly. Instead of a COM, we use the Mg ion’s position as the location for Chls, and for the Crts only the COM resulting from the atoms in the conjugated carbon chain.

The orientation factors *κ*^2^ for the chromophores and states in the CP29 model were derived from TDMs that approximately correspond to high-level quantum mechanics calculations.^7,45^ Namely, the Chl Q band TDM was associated to the Qy state and the B band TDM to the B_x_ state, aligned along the Chl atoms NB-ND and C4A-C4C, respectively. For Crts S_2_ states, TDMs were aligned along the axis connecting the C12 and C32 atoms (exception: Bcr, C12 and C19 atoms due to different atom numbering; same physical representation).

## Results and Discussion

### Theoretical values for the higher state fluorescence yields

Eqs. (4) and (5) provide the fluorescence yields listed in Table 2. The computed values coincide very well with the experimental estimates, within the same order of magnitude. To bias against the possibility of the high-energy EET processes investigated here, the lower yield will be used for the following computations, as highlighted in bold in Table 2.

**Table 2:**
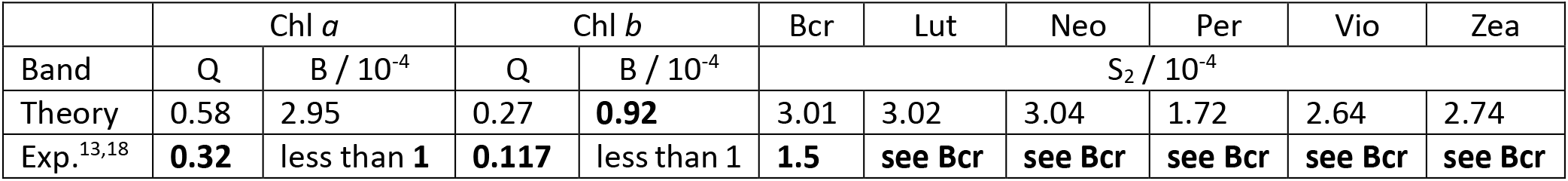
Computed and experimentally known values for the fluorescence yields of Chl and Crt states. For the Crts, the IC lifetime for Bcr (163 fs) was used in all cases. Bold values are the ones used throughout this article for calculations.

|  | Chl *a* |  | Chl *b* |  | Bcr | Lut | Neo | Per | Vio | Zea |
| --- | --- | --- | --- | --- | --- | --- | --- | --- | --- | --- |
| Band | Q | B / $10^{-4}$ | Q | B / $10^{-4}$ | $S_2 / 10^{-4}$ | | | | | |
| Theory | 0.58 | 2.95 | 0.27 | **0.92** | 3.01 | 3.02 | 3.04 | 1.72 | 2.64 | 2.74 |
| Exp.<sup>13,18</sup> | **0.32** | less than 1 | **0.117** | less than 1 | **1.5** | see Bcr | see Bcr | see Bcr | see Bcr | see Bcr |

### Q-Q, B-B and B → S_2_ FRET parameters

The computed Chl-Chl parameters are listed in Table S4 of the SI. For the Q-Q homotransfer in Chl *a* and *b*, *J* (Eq. (2)) is computed to be 55.1 and 41.7 (in units of 10^14^ M^-1^ cm^-1^ nm^4^), respectively. For the B-B homotransfer, we can expect a lower overlap, due to the scaling with *λ*^4^, and indeed the values are 12.0 for Chl *a* and 24.0 for Chl *b*. When including the full Chl absorption spectra, allowing for B → Q EET, these values increase to 12.1 and 24.4, showing that this interaction can be safely neglected (data not shown). The resulting R0 Q-Q and B-B values can be found in Figure 2A, and Table S4 of the SI. The model predicts the radii for the Q-Q homotransfer correctly (computed up to 73.8 vs. experiment 50-75 Å for Chl *a*).^46^ Using the same approach for the higher energy B-B transfer (blue columns in Figure 2A), drastically shorter R0 values are obtained (down to about 1/7 of Q-Q, *e.g.*, 11.1/73.8 for Chl *a*), which would superficially indicate that B-B EET may be less relevant than Q-Q EET. This, however, only holds when considering individual pairs of donors/acceptors. But before investigating a real system, such as CP29, the FRET parameters for Chl→Crt EET need to be introduced first.

**Figure 2:**
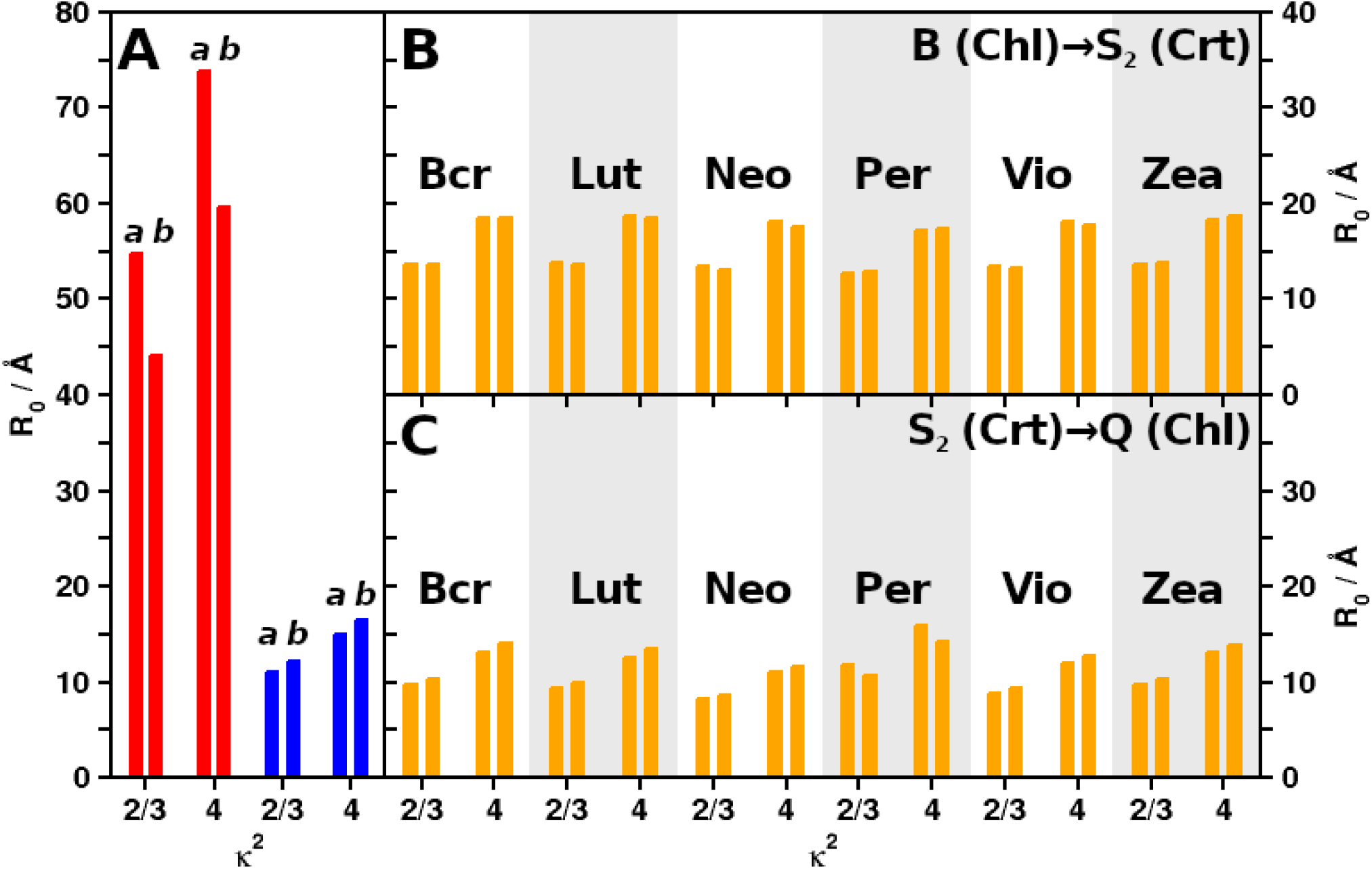
R_0_ for different values of *κ*^2^ and Chls a/b (left/right in each column pair, only indicated in (A)). (A) Chl-Chl homotransfer (red: Q-Q. blue: B-B). (B) For FRET from B (Chls) to S_2_ (Crts). (C) For FRET from S_2_ (Crts) to Q (Chls). Fluorescence yields used as listed in Table 2.

For the Chl→Crt (B→S_2_) EET, the corresponding values can be found in Figure 2B (see Table S5 in the SI for detailed values). Per as acceptor is only shown for completeness: it is well known that Per is the main light harvesting pigment in PCP and acts primarily as a donor, not an acceptor.^2,3,32,38,39^ Regardless, B→S_2_ EET exhibits R0 of about 13 to 18 Å. The listed values do not differ qualitatively (for a given *κ*^2^), although Neo and Vio exhibit a slight preference for Chl *a* as donor. The strongest preference shows Zea (for Chl *b*). Since Vio is replaced by Zea in light-harvesting complex II (LHCII) under light stress,^20,34,47^ the different affinities for Chl *a* or *b* to Vio and Zea might be relevant in this context (see also later in this article). Like for the B-B EET, the immediate impression is deceiving: The small differences in R0 could suggest that the relative coupling differences are small. However, it should be remembered that the EET rates in FRET correlate with 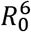, which greatly increases even minute differences.

To pursue this argument, the ratio between the EET rates of two acceptors coupled to the same donor should be considered. The relative efficiency is then

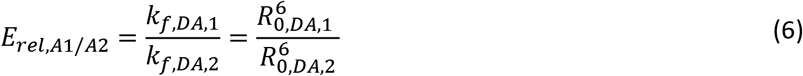

if *r*_*DA*,1_ = *r*_*DA*,2_. In peripheral light harvesting complexes, Lut is the most prevalent Crt.^35^ Considering a competing situation between Lut and Chl *a*, *E_rel,Lut/Chl a_* would become 13.8^6^/11.1^6^ = 3.69 (for *κ^2^ = 2/3*). This means that a small difference of 2.7 Å in R0, as depicted in Figure 2A and Figure 2B for B → B/S_2_ (Chl *a*/Lut), already results in a nearly fourfold preference for EET to Lut for a weakly coupled chromophore pair.

### FRET in the B band region of CP29

While the smaller B-B and B→S_2_ R0 shown in Figure 2 do not allow for long-range EET like in the Chl Q band (*R*_0,*Chl a/Chl a,Q-Q*_ > 55 Å^24^), tightly packed assemblies such as photosynthetic antennae may also exhibit FRET for processes with smaller R0. To illustrate an actual B band FRET network, CP29^35^ will be used as example, including all B and S_2_ interactions (Figure 3, details in Table S6 of the SI). The displayed ratios between IC (grey) and the two classes of EET processes (to Chls, blue, or to Crts, orange) clearly supports two core notions: (i) B-B EET exists and mostly outcompetes B → Q IC. Only the Chl 614 shows an IC larger than the sum of the available B-B processes. B-B EET would make up 77.7% of all Chl B band processes, were it not for the presence of Crts. Due to the small R0, this can only be explained by either very close distances between individual pairs or simply the vast number of potential acceptors. It turns out that it is actually a mix of both, as individual Chls exhibit different environments. (ii) A large fraction (39.1% on average) of Chl *a/b* B band excitations is preferentially transferred to Crts. With Crts, the B-B EET fraction is only 47.3% instead of 77.7%. Especially Chl 602, 603, 608, 610 and 612 would otherwise preferentially perform B-B EET, as the IC ratio is comparatively weak. Some other Chls show nearly a half/half B/S_2_ EET distribution (606, 611) or retain a preference for B-B EET (604, 607, 609, 613, 615, 616). 614 keeps its IC preference also in the presence of Crts.

**Figure 3:**
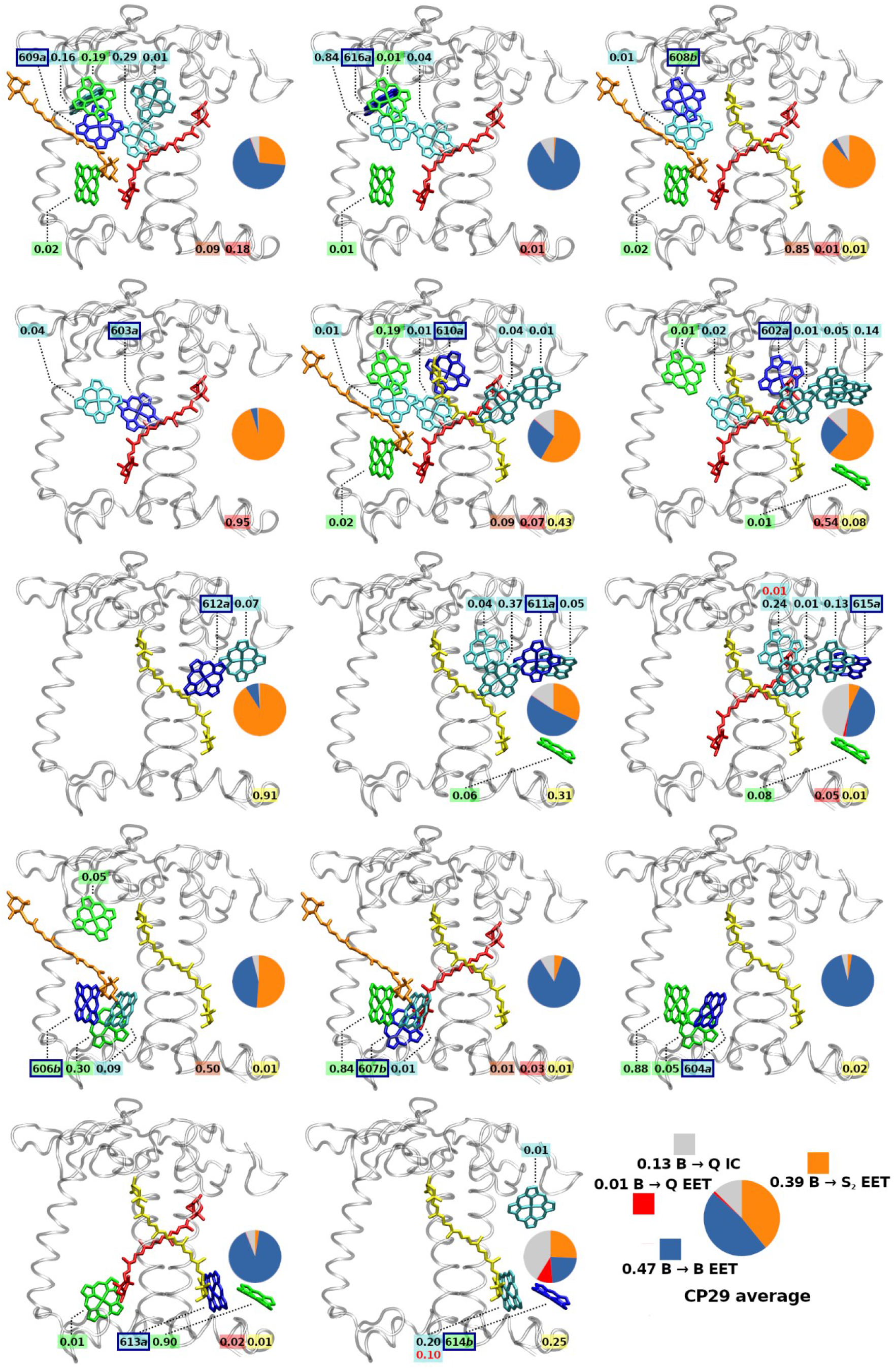
Acceptor pigments (rate fraction larger than 0.005) for FRET B → X processes of B band excited Chl donors, shown for each Chl in CP29 (donor molecule in dark blue). Acceptor Chls a in cyan, Chl b in green and Crts in red/orange/yellow (Neo/Vio/Lut). E_FRET,total_ for each acceptor from the current donor shown as black (B → B and B → S_2_) or red (B → Q) numbers. Circles: Rate fraction of B → Q internal conversion (IC, grey) vs. all FRET B → Q (red), B → B (blue) and B → S_2_ EET processes (orange)..

Regarding notion (i), some Chls, namely 603, 608, 610 and 612, donate almost exclusively to Crts. These high efficiencies imply a close distance to a specific Crt (*e.g.*, for Chl 612: Lut, 6.6 Å). For such cases, PDA is probably not sufficient anymore, and the effect might be exaggerated. For Chl 614, the immediate environment apparently does not contain many potential acceptors. This leads to the observed IC prevalence, which in turn also seemingly exaggerates the transfer to the Q band (Figure 3, Chl 614 subgraph). This is yet only due to model limitations, as CP29 itself is normally part of the photosystem II (PSII) supercomplex. The comparatively low B band EET performance of Chl 614 is therefore due to the absence of the adjacent antennae LHCII and CP24. *E.g*., including the Bcr of the neighboring CP24 complex in a test calculation (using the parameters for the CP29 Neo binding pocket) increases the fraction of CP29 Chl→Crt EET from 39.1% to 40.2%, just by adding a single additional acceptor. While there are also no immediately obvious differences between Chl *a* or *b* visible in Figure 3, we were able to identify a new specific role for this pigment type; this will however be the topic of the next article in this series to remain focused on the Crts here.

It must be noted that the Chl→Crt process comes after an optical filter effect: The strong overlap in absorption spectra (not to be confused with *J*) yields a direct competition between Chls and Crts for any photon that can be absorbed by both B and S_2_ states. This enhances the effective B band depletion by Crts, as it does not matter if a photon is directly absorbed by Crts or if the S_2_ excitation is resulting from B→S_2_ EET; the result is the same. This aspect will also be explored in the next article in this series.

Concluding this section, Crts drastically reduce Chl→Chl B-B FRET processes. This role is facilitated twofold, first by direct competition with Chl excitations (a filter effect^48^), and second, by competitively depleting Chl B band excitations via EET. The EET competition results in a reduction of Chl B-B EET by a factor of 1-(47.3/77.7) = 0.39 in an isolated CP29 complex. This role explains their ubiquitous presence in the antennae and RCs, preventing EET of potentially harmful^15–17^ B band excitations to the RCs. Further details on this crucial role will be provided at the end of this article.

### S_2_ → Chl FRET

An important issue in discussing the potential of back-transfer to Chls from S_2_ of Crts is the energetic location of the Crt S_2_ state. The emission spectrum of Crt S_2_ is experimentally and theoretically known (Table 1).^7^ It should however be noted that the experimental 0-0 excitation energies of Crts, typically listed at around 500 nm for the Crts discussed here,^1^ do not correspond to the FRET-relevant emission energies. Instead, the overall emission spectrum is relevant, not the energetic location of individual Franck-Condon factors such as for the 0-0 transition or a maximum intensity. While the adiabatic transition may be important for the photophysical characterization of the Crts, the mean of the Crt emission spectra is not at all located close to 0-0.^7^ The general depiction of S_2_ being energetically far from the Chl Q states is thus very misleading in terms of Crt-Chl FRET capabilities.

Keeping the above in mind, the FRET parameters for a Crt → Chl EET process, from Crt S_2_ to Chl, can be found in Table S7 of the SI., R0 values are shown in Figure 2C. Table S7 of the SI also shows that Crt emission in almost all cases preferentially couples to the Q band, not B (the exception being Neo → Chl *b*, for which the overlap is evenly shared between B and Q).

All interactions for a Crt-Chl FRET indicate a preference for EET to Chl *b* instead of Chl *a*, apart from Per, which is (unsurprisingly^49^) preferring Chl *a* as acceptor. The model thus correctly predicts the corresponding biological function of Per. Furthermore, Per is much better suited for back-transfer EET to Chls in general, as shown by the much stronger spectral overlap (from SI: 12.0/6.2, Chl *a/b*), as compared to the next strongest donor Crt, Bcr (from SI: 3.7/5.5). Per, thus, has likely evolved from a primarily B band-depleting pigment to a primarily Q donating Crt. Among the other Crts, Bcr and Zea are expectedly similar, as they are structurally almost identical. The donor ability of the Crts, excluding Per, depends on the length of their conjugated system, which can be easily derived from a particle-in-a-box perspective (short length → higher energies). The resulting order Bcr/Zea → Lut → Vio → Neo applies consistently for the spectra.

From comparing Figure 2B and Figure 2C, it can be inferred that the back-transfer from Crts should be much less efficient (in a PDA/FRET model) than the EET from the B band to Crts, due to the smaller *R*_0,*Crt/Chl*_ values. However, now with Chls as the acceptors, the radius can be much smaller due to Chls being more numerous. The corresponding results for a CP29 network with Crts as donors can be found in Figure 4 (and Table S8 in the SI).

**Figure 4:**
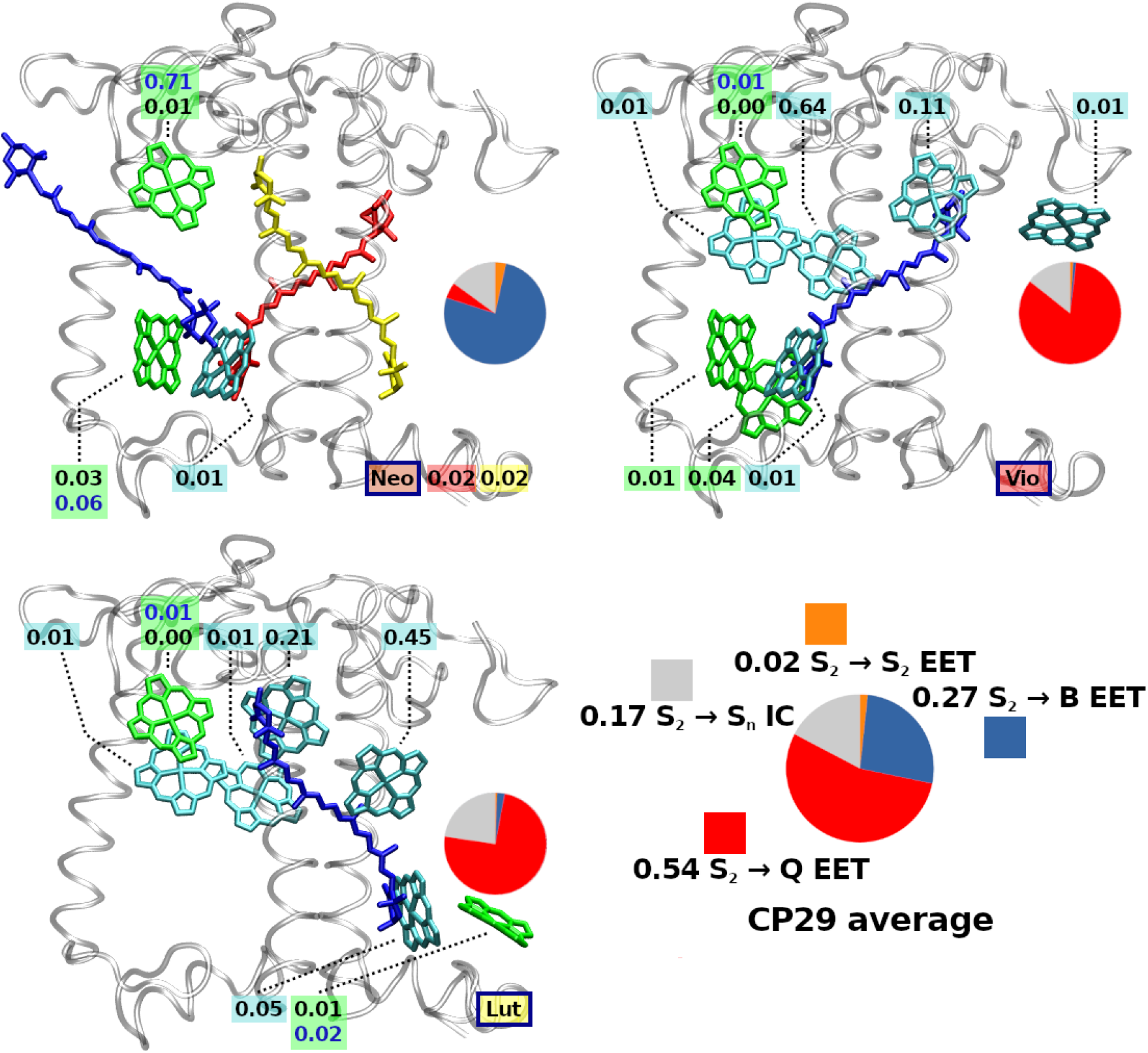
Acceptor pigments (rate fraction larger than 0.005) for FRET S_2_ → X processes of S_2_ excited Crt donors, shown for each Crt in CP29 (donor molecules in blue). Acceptor Chls a in cyan, Chls b in green and Crts in red/yellow (Vio/Lut). E_FRET,total_ for each acceptor from the current donor shown as black (S_2_ → Q and S_2_ → S_2_) or blue (S_2_ → B) numbers. Circles: Rate fraction of S_2_ → S_n_ internal conversion (IC, grey) vs. all FRET S_2_ → Q (red), S_2_ → B (blue) and S_2_-S_2_ EET processes (orange).

The Crt back-donation shows that Lut and Vio basically exhibit the same pattern, with a slightly stronger IC to the Crt dark state (S_1_) in Lut (22.7% vs. 14.3%). In contrast, Neo is primarily coupled to the B state of Chl 608 (71%), which distinguishes this Crt from all others. This is due to the shorter conjugated chain in Neo compared to Vio or Lut, which raises the emission energies significantly. Again, it must be stressed that CP29 is not an isolated complex – especially the Neo binding site is located such that one half of the bound pigment is not inside the CP29 complex but arranged towards other complexes (LHCII in this case). It is thus likely that for a full PSII supercomplex, Neo shows a slightly different coupling pattern.

Despite the Förster radii being much smaller, back-donation from S_2_ of Crts to Chl Q states via FRET is found to be similarly efficient as the B → S_2_ FRET process. Within the PDA, they can do so by either strong individual coupling (*e.g.*, Vio-Chl 603) or by coupling to several different Chl acceptors with similar preference (*e.g.*, Lut-Chls 610/612).

### Testing the effect of Crt replacement: Zea for Vio

The above calculations were repeated for a slightly modified CP29 model, this time containing Zea instead of Vio (discussed here only in the text for sake of brevity). Although the replacement is more related to NPQ in LHCII and its specific fourth Crt binding site,^20,34,47^ the effects should allow to draw preliminary conclusions for a future LHCII model. The calculation results in S_2_ → Q donation rates of Zea being 4.1 times higher than those of Vio. At the same time, the sum of all acceptor rates went down by a factor of 0.36. Overall coupling patterns (*i.e.*, preferential FRET donors/acceptors) remain identical upon replacement of Vio by Zea in the CP29 model.

Introducing Zea changes the focus from B band coupling to Q band coupling, at least within our model. It remains to be seen if such a change is beneficial under light stress; within the present simulations, no indication is found. For the Zea-induced quenching in CP29 found by Crimi *et al*.,^50^ the model presented here is unfortunately not precise enough – the simulated Crts do not differ in *τ*_*S*_2_→*S_n_*_.

## Summary and conclusions

It is shown that EET is not only occurring between the Chl Q band states. Instead, all states with significant TDM participate in EET, even those that undergo rapid IC. This results in Chl B-B EET, which might have detrimental consequences.^15–17^

It is also shown that a simple, efficient solution to the Chl B-B EET problem is the presence of Crts. Crts efficiently compete for Chl B state excitations, both via initial spectral filtering (discussed in detail in the next article of this series) and then through a high likelihood of acting as FRET acceptors (in CP29, at least 39% B-B reduction). Due to rapid internal conversion, and their strongly shifted emission spectrum, they cannot donate the energy back into the B band. We have shown that this population can instead efficiently be donated (by Lut, Vio) into the Chl Q band or re-routed (Neo) to other Chl B bands.

Crts thus provide an important, but so far overlooked, function in photosynthetic light harvesting: re-routing of blue excitations and, as result, reducing the B-B EET between Chls.

### Implications

From an evolutionary perspective, it is interesting to note that even the LHC predecessors, (one-)helical membrane proteins, contain Crt binding sites.^51,52^ On the other hand, the problem of B-B EET is definitely distance-dependent (due to the comparatively small R0 values). It seems, that in earlier evolutionary stages, before Chls became clustered, B-B EET may have been simply defused via IC. Crts or other remedies to the B-B EET would only be required to overcome a certain Chl density limit.

There are, however, alternative solutions to the B-B EET problem. One might be simply a high-light avoiding movement of the organism, but higher plants lack this ability. Another one would be a protein-based re-routing mechnism.^44^ There might be other, related possibilities: For example, bacteriochlorophylls appear to be potential candidates as well, since their B band absorption may be less.^53^ However, it is commonly assumed that BChls have evolved as a response to other Chls, harvesting light in spectral regions to avoid competition by Chls.^54,55^

Finally, there is the explicit possibility of Chl *b* playing a role in the B-B EET context. The next article in this series will aim to shed light on a potential new function of Chl *b* as a photoprotective pigment.

## Supporting information

Supplementary material (all)

## Acknowledgements

J.P.G. gratefully acknowledges funding by the Deutsche Forschungsgemeinschaft (DFG), grant number 393271229. H.L. gratefully acknowledges funding by the Czech Science Foundation, GAČR (grant no. 22-17333S). The authors thank Prof. Dr. Peter Saalfrank for his valuable comments on an earlier version of the manuscript.

## Author contributions

J.P.G. provided the theoretical concept, calculations, and figures, as well as the initial draft. H.L. provided conceptual guidance, the experimental spectra and edited the draft. Both authors contributed equally to the finalization of the manuscript.

