## Supplementary material (all) for "Excitation energy transfer between higher excited states of photosynthetic pigments: 1. Carotenoids facilitate B → Q band conversion in chlorophylls"

**Supporting Information**

Jan P. Götze[1,]* and Heiko Lokstein[2]

[1]*Institut für Chemie und Biochemie, Freie Universität Berlin, Arnimallee 22, 14195 Berlin, Germany; ORCID ID 0000-0003-2211-2057*

[2]*Department of Chemical Physics and Optics, Charles University, Ke Karlovu 3, 121 16 Prague, Czech Republic; ORCID ID 0000-0001-6739-4612*

Table of contents:

- CP29 site energies
- CP29 distance matrices
- Chl donor FRET parameters
- CP29 summed rates
- Crt-Chl FRET parameters
- CP29 back-donation summed rates

CP29 site energies

*Table S1: $Q_y$ and $B_X$ energy shifts in CP29 derived from data in Petry and Götze, relative to averages of Chl a or b states/sites.*

| **Site ID** | **602*a*** | **603*a*** | **604*a*** | **606*b*** | **607*b*** | **608*b*** | **609*a*** |
|---|---|---|---|---|---|---|---|
| **Q shift / meV** | -11 | +1 | +7 | +17 | -8 | +8 | +21 |
| **B shift / meV** | -32 | +22 | -1 | +40 | +55 | -54 | +19 |
| **Site ID** | **610*a*** | **611*a*** | **612*a*** | **613*a*** | **614*b*** | **615*a*** | **616*a*** |
| **Q shift / meV** | +3 | -24 | +18 | -1 | -17 | -51 | +40 |
| **B shift / meV** | -4 | -65 | +15 | +32 | -43 | -76 | +93 |

CP29 distance matrices

*Table S2: CP29 Chl-Chl distance matrix, according to chain R of PDB entry 5XNL, Mg-Mg distances, in Å*

| 603 | 604 | 606 | 607 | 608 | 609 | 610 | 611 | 612 | 613 | 614 | 615 | 616 | |
|---|---|---|---|---|---|---|---|---|---|---|---|---|---|
| 11.7 | 24.9 | 26.8 | 24.9 | 21.5 | 17.9 | 15.9 | 15.6 | 17.9 | 18.9 | 24.4 | 13.1 | 20.7 | 602 |
| | 19.3 | 18.3 | 15.1 | 17.8 | 9.3 | 18.3 | 24.5 | 22.8 | 19.6 | 28.1 | 22.9 | 13.0 | 603 |
| | | 7.9 | 11.7 | 18.2 | 17.4 | 18.3 | 26.8 | 19.2 | 20.6 | 26.6 | 32.0 | 27.0 | 604 |
| | | | 9.5 | 15.9 | 13.7 | 20.8 | 32.2 | 25.2 | 25.8 | 33.1 | 36.1 | 22.5 | 606 |
| | | | | 22.0 | 15.3 | 24.8 | 32.0 | 26.9 | 20.8 | 29.2 | 33.0 | 22.3 | 607 |
| | | | | | 10.4 | 11.5 | 27.9 | 22.2 | 30.8 | 37.2 | 33.1 | 17.2 | 608 |
| | | | | | | 16.5 | 28.6 | 24.6 | 26.2 | 34.3 | 30.1 | 9.82 | 609 |
| | | | | | | | 16.9 | 11.7 | 24.6 | 29.0 | 24.4 | 23.7 | 610 |
| | | | | | | | | 9.2 | 18.6 | 18.0 | 12.3 | 34.5 | 611 |
| | | | | | | | | | 18.3 | 19.4 | 19.7 | 32.4 | 612 |
| | | | | | | | | | | 9.2 | 17.3 | 32.6 | 613 |
| | | | | | | | | | | | 18.1 | 40.9 | 614 |
| | | | | | | | | | | | | 33.4 | 615 |

*Table S3: CP29 Chl-Crt distance matrix, according to chain R of PDB entry 5XNL, Mg-Crt(COM) distances, in Å. Crt(COM) (center of mass) computed for conjugated carbon atoms in the Crt chain.*

| | 602 | 603 | 604 | 606 | 607 | 608 | 609 | 610 | 611 | 612 | 613 | 614 | 615 | 616 |
|---|---|---|---|---|---|---|---|---|---|---|---|---|---|---|
| Lut | 16.7 | 18.1 | 13.0 | 18.6 | 20.6 | 17.7 | 19.1 | 9.7 | 14.2 | 6.6 | 16.6 | 20.5 | 21.8 | 27.7 |
| Neo | 29.2 | 23.8 | 14.9 | 12.2 | 21.3 | 10.0 | 16.3 | 17.3 | 32.8 | 25.1 | 33.2 | 39.2 | 39.5 | 24.4 |
| Zea | 11.2 | 9.6 | 15.7 | 18.8 | 15.5 | 20.6 | 15.6 | 16.3 | 17.6 | 15.6 | 11.0 | 18.9 | 17.7 | 22.2 |

Chl donor FRET parameters

*Table S4: FRET parameters computed for Q-Q and B-B homotransfer in Chls a and b. J in $10^{14}$ $M^{-1}$ $cm^{-1}$ $nm^4$, $R_0$ in Å.*

| | $J_{Chl\,a/Chl\,a}$ | $J_{Chl\,b/Chl\,b}$ | $R_{0,Chl\,a/Chl\,a}$ | | $R_{0,Chl\,b/Chl\,b}$ | |
|---|---|---|---|---|---|---|
| $\kappa^2$ | / | / | 2/3 | 4 | 2/3 | 4 |
| **Q band** | 55.1 | 41.7 | 54.7 | 73.8 | 44.2 | 59.6 |
| **B band** | 12.0 | 24.0 | 11.1 | 14.9 | 12.2 | 16.5 |

*Table S5: FRET parameters computed for a Soret Chl a/b to Crt FRET process. J in $10^{14}$ $M^{-1}$ $cm^{-1}$ $nm^4$, $R_0$ in Å.*

| | **Chl-to-Crt** | $J_{Chl\,a/Crt}$ | $J_{Chl\,b/Crt}$ | $R_{0,Chl\,a/Crt}$ | | $R_{0,Chl\,b/Crt}$ | |
|---|---|---|---|---|---|---|---|
| | $\kappa^2$ | / | / | 2/3 | 4 | 2/3 | 4 |
| **Carotenoid** | **Bcr** | 42.79 | 46.12 | 13.7 | 18.4 | 13.6 | 18.4 |
| | **Lut** | 45.37 | 47.06 | 13.8 | 18.6 | 13.7 | 18.5 |
| | **Neo** | 39.45 | 33.85 | 13.5 | 18.2 | 13.0 | 17.5 |
| | **Per** | 27.34 | 33.11 | 12.7 | 17.1 | 12.9 | 17.4 |
| | **Vio** | 40.34 | 37.58 | 13.5 | 18.2 | 13.2 | 17.8 |
| | **Zea** | 40.87 | 48.75 | 13.6 | 18.3 | 13.8 | 18.6 |

CP29 summed rates

*Table S6: IC rates (in $ps^{-1}$) for Chl B band donation, ratios to the sum of all processes.*

| Chl | IC | IC Fraction | Q Fraction | B Fraction | Crt Fraction |
|---|---|---|---|---|---|
| **602** | 10 | 0.13 | 0.00 | 0.25 | 0.62 |
| **603** | 10 | 0.01 | 0.00 | 0.04 | 0.95 |
| **604** | 10 | 0.04 | 0.00 | 0.93 | 0.03 |
| **606** | 17.2 | 0.04 | 0.00 | 0.45 | 0.51 |
| **607** | 17.2 | 0.09 | 0.00 | 0.86 | 0.05 |
| **608** | 17.2 | 0.08 | 0.00 | 0.04 | 0.88 |
| **609** | 10 | 0.05 | 0.00 | 0.68 | 0.27 |
| **610** | 10 | 0.14 | 0.00 | 0.28 | 0.58 |
| **611** | 10 | 0.15 | 0.00 | 0.52 | 0.32 |
| **612** | 10 | 0.01 | 0.00 | 0.08 | 0.91 |
| **613** | 10 | 0.06 | 0.00 | 0.91 | 0.02 |
| **614** | 17.2 | 0.40 | 0.10 | 0.22 | 0.28 |
| **615** | 10 | 0.46 | 0.02 | 0.45 | 0.07 |
| **616** | 10 | 0.09 | 0.00 | 0.90 | 0.01 |

### Crt-Chl FRET parameters

*Table S7: FRET parameters computed for a Crt bright emission ($S_2$) to Chl FRET process. Q band only (FRET to full spectrum in parentheses). $J$ in $10^{14}$ $M^{-1}$ $cm^{-1}$ $nm^4$, $R_0$ in Å. Neoxanthin emission unavailable, using octaene instead.*

| | Chl-to-Crt | $J_{Crt/Chl\,a}$ | $J_{Crt/Chl\,b}$ | $R_{0,Crt/Chl\,a}$ | | $R_{0,Crt/Chl\,b}$ | |
|---|---|---|---|---|---|---|---|
| | $\kappa^2$ | / | / | 2/3 | 4 | 2/3 | 4 |
| Carotenoid | **Bcr** | 3.72 (3.72) | 5.51 (5.73) | 9.7 | 13.1 | 10.4 | 14.0 |
| | **Lut** | 2.92 (2.93) | 4.29 (5.35) | 9.3 | 12.6 | 10.0 | 13.4 |
| | **Neo** | 1.38 (1.38) | 1.78 (3.59) | 8.3 | 11.1 | 8.6 | 11.6 |
| | **Per** | 12.02 (12.02) | 6.15 (6.15) | 11.8 | 16.0 | 10.6 | 14.3 |
| | **Vio** | 2.13 (2.14) | 3.10 (4.73) | 8.9 | 12.0 | 9.4 | 12.7 |
| | **Zea** | 3.66 (3.66) | 5.22 (5.66) | 9.7 | 13.1 | 10.3 | 13.9 |

CP29 back-donation summed rates

*Table S8: IC rates (in $ps^{-1}$) for Crt $S_2$ donation, ratios to the sum of all processes.*

| Crt | IC | IC Fraction | Q Fraction | B Fraction | Crt Fraction |
|---|---|---|---|---|---|
| **Lut** | 6.135 | 0.23 | 0.74 | 0.02 | 0.01 |
| **Vio** | 6.135 | 0.14 | 0.84 | 0.01 | 0.01 |
| **Neo** | 6.135 | 0.15 | 0.05 | 0.76 | 0.03 |
